# Coordinate-Dependent Neuroinflammation and Metabolic Disruption Following Experimental Traumatic Brain Injury

**DOI:** 10.1101/2025.08.05.668726

**Authors:** Shruti Kaza, Dilmareth Natera-Rodriguez, Andrew Grande

## Abstract

Traumatic brain injury (TBI) is a leading cause of long-term neurological dysfunction, with secondary neuroinflammation playing a pivotal role in disease progression. Microglia, the brain’s resident immune cells, respond to injury in a spectrum of activation states, influencing neuronal survival and cognitive recovery. While past studies have broadly examined neuroinflammatory responses following TBI, the influence of injury approach—specifically, anterior vs. posterior impact—on microglial activation and neuronal damage remains unexplored. This study investigated how the site of controlled cortical impact (CCI) in a moderate TBI model alters neuroinflammatory responses and neuronal vulnerability, providing key insights into coordinate-specific mechanisms of injury.

## 1. Introduction

Head trauma is responsible for over 10% of deaths globally, despite advancements in medical care^28^. In the United States, 3.5 million cases of traumatic brain injury (TBI) occur annually, making it a significant factor in fatal outcomes associated with nervous system damage. Understanding the physiological changes in the brain following moderate TBI (mTBI) is essential^28^. mTBI often results from concussions incurred in contact sports, military combat, or accidents. TBI is defined as an alteration in brain function caused by external force. Current scientific hypotheses identify two primary mechanisms for TBI: the mechanical impact of trauma and a subsequent inflammatory response that leads to dysregulated cytokine expression and proinflammatory microglial activation, ultimately resulting in neurodegeneration and cell death^27^. Investigating the cellular and molecular mechanisms underlying moderate TBI has been challenging due to the absence of standardized procedures for inducing TBI in preclinical models.

The inflammatory pathway following TBI can contribute to cognitive disorders like dementia and exacerbate the effects of neuroinflammation. Research indicates that injury triggers pathological responses that alter the intracellular dynamics of neurons^28^. This suggests that certain brain regions may be more vulnerable to neuronal damage. Thus, it is necessary to examine the effects of neuronal damage across various brain regions at the cellular level. Previous studies in mice have focused on behavioral changes post-mTBI across different brain coordinates but have not investigated variations in cellular pathways^12^.

This study diverges from earlier literature by emphasizing whole-brain analysis and coordinate-specific patterns. The anterior and posterior brain coordinates, in particular, differ in their cellular composition, biomechanical properties, and susceptibility to injury-induced changes. This is crucial for developing targeted therapies. Previous studies of inflammation with regional variation determined region-specific inflammation but through lipopolysaccharide (LPS) induced inflammation. This study does not reflect TBI-induced inflammation, identifying a crucial gap in understanding the injury-induced deficit for therapeutic interventions^10^. This study explores these regional differences by analyzing microglial activation, astrocyte proliferation, and mitochondrial dysfunction, aiming to uncover coordinate-specific insights^3^.

Following injury, activated microglia, the primary immune cells in the brain, are integral to the neuroinflammatory response. However, extended activation after TBI contributes to neurodegenerative processes^3^. Thus, it is vital to explore how microglial density and activation differ across brain regions following TBI. Microglial cells are highly dynamic and exhibit two phenotypes: proinflammatory (M1) and anti-inflammatory (M2). However, recent research suggests that these phenotypes are not strictly categorized, and both may be present post-injury. Therefore, further investigation into the role of microglia in post-injury inflammation is needed^19^. Since microglial activation and mitochondrial function are crucial indicators of neuroinflammation, exploring their variation across brain coordinates is key to identifying targeted interventions.

Additionally, neuroinflammation post-injury leads to mitochondrial dysfunction, which impacts energy metabolism. The brain utilizes almost 20% of total body energy and proper mitochondrial function is critical for meeting neuronal energy demand. Mitochondrial damage is associated with neuronal injury and cell death, compromising adenosine triphosphate (ATP) production and leading to energy deficits^22^. Features such as oxidative stress and inflammation related to neurological diseases have been linked to mitochondrial impairment.

Current research on mitochondrial dysfunction in anterior and posterior brain regions has primarily centered on Parkinson’s disease, highlighting the need for studies examining TBI-related dysfunction^19^. Previous work, such as Fischer et al. (2016), has found that TBI reduces mitochondrial respiration, increases inflammation, and triggers apoptosis^7^. Mitochondrial activity is divided between fission and fusion to meet energy demands^22^. However, there is currently a lack of research focusing on fusion-specific dynamics of mitochondria post-TBI, and analyzing mitochondrial function through a coordinate-specific lens is crucial for fostering neuronal protection and enhancing cognitive outcomes post-TBI. Hence, this study aims to investigate how anterior-posterior brain coordinates influence microglial activation and mitochondrial dysfunction following mTBI.

## 2. Hypothesis and Expected Outcomes

This study hypothesizes that microglial activation and neuroinflammation will differ significantly between anterior and posterior brain coordinates following moderate traumatic brain injury (TBI). We expect these differences to correlate with varying levels of disruption, neurotoxic microglial activation, and inflammation in their respective areas. Additionally, control samples are anticipated to show regional variations in baseline microglial activity, reflecting inherent differences in susceptibility to injury. We aim to establish a correlation between baseline neuroinflammation and subsequent mitochondrial dysfunction.

To test this hypothesis, the study encompasses five main aims:

1. Establish a Standardized Protocol for Moderate TBI in Preclinical Models

a. This aim focuses on developing a reproducible method for inducing moderate TBI using the CCI model, targeting both anterior and posterior brain regions. A standardized procedure will ensure consistent results and facilitate comparisons across studies.
2. Compare Microglial Activation Between Anterior and Posterior Brain Coordinates Following TBI.

a. This aim analyzes spatial differences in microglial density, activation states, and neuroinflammatory markers in both regions.
3. Evaluate the Relationship Between Microglial Activation and Neuroinflammation in Coordinate-Specific Injury Responses.

a. This aim seeks to understand how differences in microglial activation contribute to mitochondrial dysfunction, axonal damage, and inflammatory cytokine production in the anterior and posterior coordinates.
4. Provide Insights into Coordinate-Specific Therapeutic Strategies for TBI.

a. This objective aims to facilitate the development of targeted therapeutics for more effective treatment, recognizing that each injury is unique.
5. Analyze Mitochondrial Dysfunction Post-TBI to Identify Suitable Function-Specific Analysis.

a. This aim involves a comparative analysis of mitochondrial dysfunction after TBI to determine function-specific activity post-TBI.

## 7. Methods and Approach

All procedures adhere to protocols approved by the Institutional Animal Care and Use Committee (IACUC). TBI surgeries were performed by trained and approved lab members, following ethical guidelines and standards of care to ensure minimal harm and consistent experimental reproducibility.

Moderate TBI was induced using the controlled cortical impact (CCI) model to target anterior and posterior coordinates validated by the Allen Brain Atlas. These coordinates were chosen due to distinct cellular compositions and biomechanical properties^2^. Quantitative analysis focused on neuroinflammatory markers (Iba1, GFAP) and neuronal integrity (NeuN), with comparisons between anterior and posterior brain regions.

### 7.1 Animal model

The experimental design (Figure 1) involved C57BL/6 mice, which were divided into four groups to evaluate region-specific injury responses. The groups included: anterior TBI, posterior TBI, anterior sham (control), and posterior sham. Each group consisted of 8 mice, with evaluations performed at 7 days post-injury (DPI) to assess neuroinflammatory and neuronal changes through immunohistochemical analysis of Iba1, GFAP, and NeuN markers. This experimental design allowed for the direct comparison of inflammatory and neuronal responses between anterior and posterior brain regions while controlling for surgical effects.

**Figure 1:**
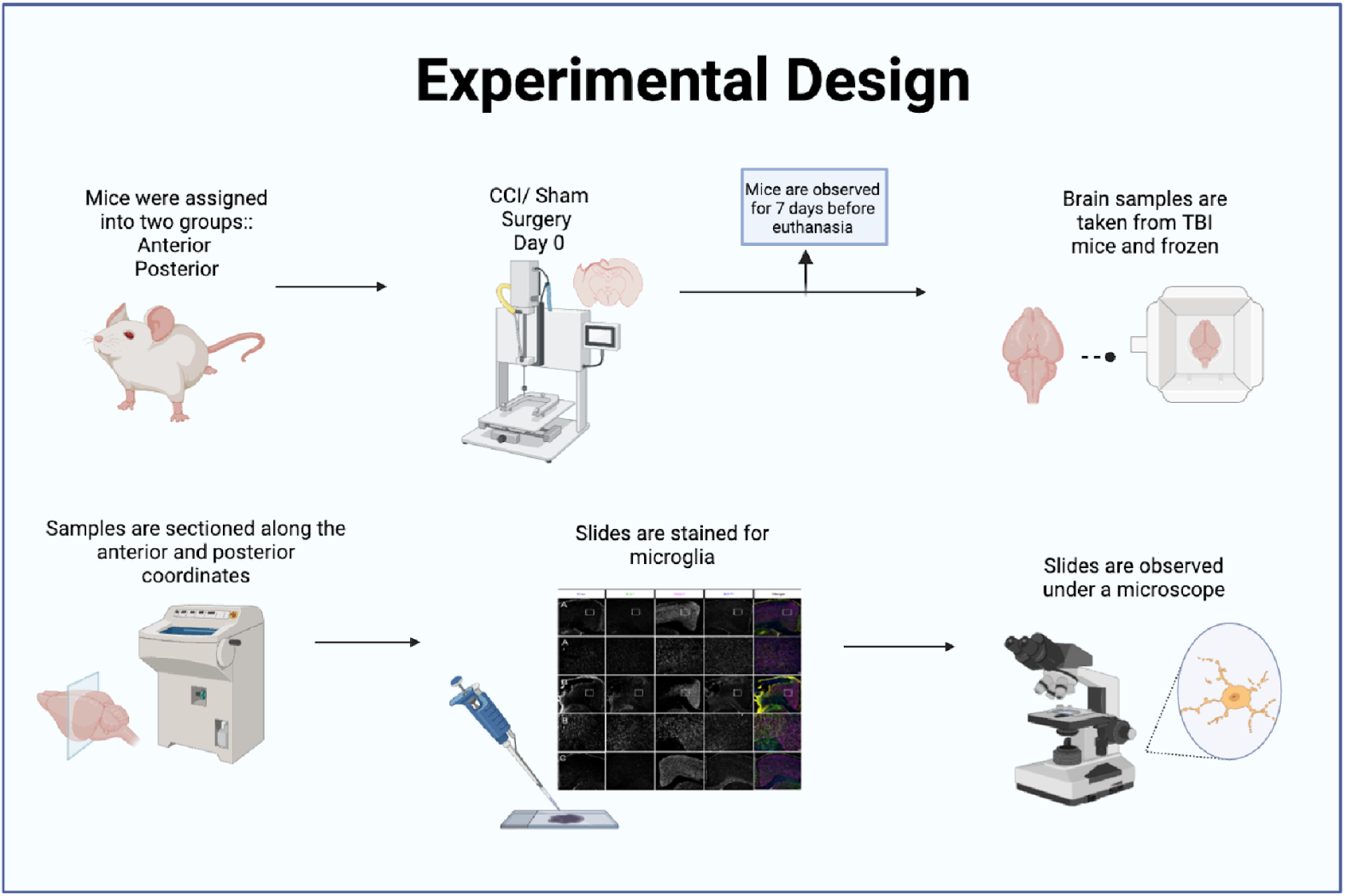
**Experimental design** of methods from CCI to IHC staining. Image created by author with biorender.com

### 7.2 Controlled Cortical Impact Traumatic Brain Injury

Moderate TBI was induced using the controlled cortical impact model (CCI) with an electromagnetic stereotaxic impactor (Impact One, Leica Microsystems Inc). Mice (7–8 weeks old) were anesthetized with 2% isoflurane and secured in a stereotaxic apparatus to allow for precise targeting of the brain coordinates. The 7-8 week timeframe for mice was to reflect a developed brain to reflect changes post-TBI, This apparatus consists of a head holder to hold the mouse in place during surgery. The scalp of each animal was shaved and wiped with alcohol. An incision was made to reflect the skin and expose the skull, and a craniotomy was performed using a dental drill. Buprenorphine (2.0 mg/kg) was administered subcutaneously for extended analgesia, which refers to the use of pain relievers for the absence of pain in the mice. A midline skin incision was made to expose the bregma, and a 2.5 mm craniotomy was performed over the primary and secondary motor cortices using a steel drill bit (Dremel), The bregma is the main reference point on the skull and was used to reference the anterior and posterior coordinates. This procedure reflects reproducible methods to induce mTBI^5^. The coordinate-specific parameters to induce mTBI are listed below and a visual representation is found in Figure 3.

For the posterior coordinate, the impact site was targeted at X = 2 mm (left lateral from bregma), Y = -2 mm (posterior from bregma), Z = -1 to -1.5 mm (depth) using a 2 mm impactor tip, with a velocity of 3 m/s, dwell time of 100 ms, and at a 15° angle. At the anterior coordinate, the impact site was positioned at X = -1.5 mm (right lateral from bregma), Y = 0, Z = -1 mm (depth) with the same parameters (velocity of 3 m/s, dwell time of 100 ms, 15° angle) using a 2 mm impactor tip.

Following surgery, mice were placed in clean home cages with heating pads and monitored for three days post-injury. Animals were euthanized 7 days post-injury (DPI) for further analysis. This procedure ensures reproducibility while minimizing harm and adhering to strict ethical guidelines. The model was chosen for its precision in targeting cortical regions and its suitability for preclinical studies of moderate TBI^5^.

The controlled cortical impact (CCI) model allows for more control over factors such as time, velocity, and depth of impact, making it more useful for studying the biomechanical factors of TBI. The severity of TBI using CCI can be controlled with impact velocity. For example, in this study, both surgeries in the anterior and posterior coordinates included a consistent impactor probe velocity of 3 m/s. Xiong et al. (2013) also point out that CCI animal models of TBI have been the most reproducible to human models^27^. This makes animal data more accurate to clinical practice and eliminates external parameters to focus solely on coordinate differences. Thus, for a standardized procedure, CCI animal models of mTBI were utilized in this study.

### 7.3 Immunohistochemical Analysis

Following perfusion with cold phosphate-buffered saline (PBS) (Genesee Scientific) and 4% paraformaldehyde (PFA) (Millipore Sigma), brains were extracted and submerged in PFA overnight at 4°C. PFA is used to preserve tissue and prepare samples for staining. This process is crucial for preserving tissue morphology and protein integrity by regulating pH (a measure of acidity) and osmolarity (concentration of solute particles). After three PBS rinses, brains were equilibrated overnight at 4°C in increasing sucrose concentrations (10%, 20%, 30%), to preserve tissue structure, and embedded in Tissue-Tek OCT (Andwin Scientific) on dry ice. The frozen blocks prepared with OCT were used for cryosectioning. Frozen blocks were stored at -80°C and sectioned coronally at 25 µm thickness using a cryostat (Leica, -20°C) into six semi-serial sets (∼180 µm between sections). A sectioning thickness of 25 µm was chosen to maintain structural integrity and provide optimal penetration of stains and antibodies. Furthermore, sections that are too thin are susceptible to wrinkles and folds, affecting the integrity of results^11^. Sections were collected on Superfrost Plus slides (Fisherbrand) to prepare for staining.

To stain these samples, this study utilized fluorescent immunohistochemistry (IHC). Fluorescent IHC allows for precise detection of neuronal markers and multiple dyes with higher image resolution^26^. Sections were post-fixed in 4% PFA, rinsed with PBST (PBS + 0.1% Triton X-100), and blocked with 10% normal donkey serum in PBST for 60 minutes. Tissue fixation involves cross-linking proteins to preserve structure for further analysis done with PBS. Donkey serum was used as a blocking agent in IHC to reduce the non-specific binding of antibodies to the tissue samples, which improves the accuracy of the staining. Slides were incubated overnight at 4°C with primary antibodies targeting Iba1, GFAP, NeuN, and MFN1 (Table 1), which were chosen to evaluate inflammation, neuronal integrity, and mitochondrial activity. These antibodies will illustrate fluorescence under the microscope. After washing, secondary antibodies (with DAPI) were applied for 60 minutes at room temperature, followed by mounting with ProLong Glass Antifade Mountant (Thermo Fisher). The 60 minute period allows for sufficient antibody-antigen binding without risking excessive background non-specific binding^16^. Antifade is used to preserve fluorescence signals and prevent photobleaching where the dyes fade. This process allows for increased sensitivity and accuracy of fluorescence imaging.

**Table 1.** Immunohistochemistry Reagents and respective details that were referenced during IHC staining.

| Antigen | 1° Antibody | Vendor | Catalog No. | 2° Antibody | Vendor | Catalog No. |
| --- | --- | --- | --- | --- | --- | --- |
| Macrophage/<br>Microglia | Iba1 | Wako | 019-19741 | Donkey Anti Rabbit (IgG) Alexa Fluor 555 | Abcam | ab150074 |
| Reactive Astrocytes | GFAP | Lifespan Biosciences | LS-C7076 GFAP | Donkey Anti Rabbit (IgG) DyLight 488 | Novus | NBP1-75 278 |
| Neurons | NeuN | Abcam | ab104224 | Donkey Anti Mouse (IgG) Alexa Fluor® 647 | Abcam | ab150107 |
| Nuclei | DAPI | Thermo Scientific | 62248 | N/A | N/A | N/A |
| Mitochondria | MFN1 | Invitrogen | 5-24789 | Mouse 647 | Abcam | ab150107 |

*Note: Mitochondria analysis with MFN1 was solely conducted through qualitative assessment of immunofluorescence.*

**Table 2.**
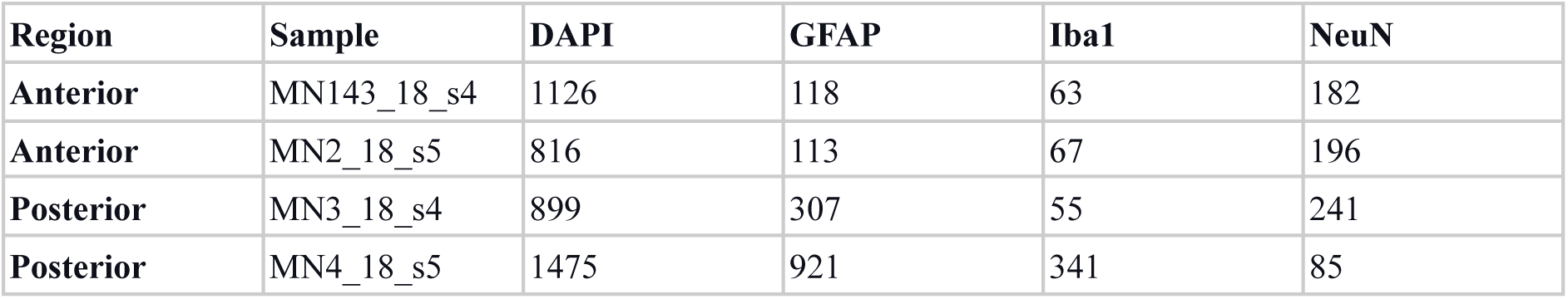
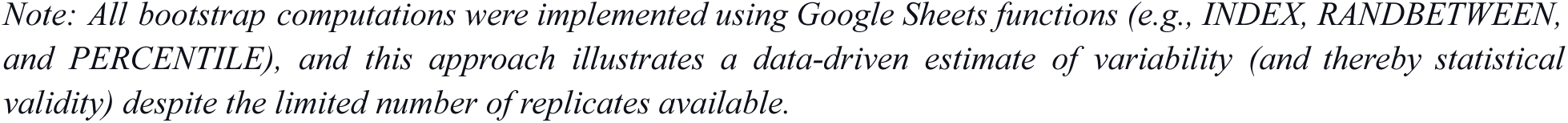
**Quantitative Microglial Density** representing raw data from cell quantification via QuPath software.

| Region | Sample | DAPI | GFAP | Iba1 | NeuN |
| --- | --- | --- | --- | --- | --- |
| Anterior | MN143_18_s4 | 1126 | 118 | 63 | 182 |
| Anterior | MN2_18_s5 | 816 | 113 | 67 | 196 |
| Posterior | MN3_18_s4 | 899 | 307 | 55 | 241 |
| Posterior | MN4_18_s5 | 1475 | 921 | 341 | 85 |

*Note: All bootstrap computations were implemented using Google Sheets functions (e.g., INDEX, RANDBETWEEN, and PERCENTILE), and this approach illustrates a data-driven estimate of variability (and thereby statistical validity) despite the limited number of replicates available.*

Images were captured using a Leica DMi8 microscope and processed in ImageJ Fiji (v1.53q) software. Microglial activation was analyzed via Iba1 staining, with GFAP staining for astrocytes to study cell structure and spatial distribution. High-resolution imaging was used to map mitochondrial and microglial changes across anterior and posterior coordinates^6^.

The IHC process allows for qualitative assessment and cell quantification of the anterior versus posterior results. This gathers the data necessary for completing coordinate and functional analysis per the study’s aims.

### 7.4 Cell Quantification

Cell quantification was performed using QuPath (v0.3.2) software to analyze IHC-stained brain sections. Cell quantification through QuPath is a method to measure fluorescence intensity or quantify and identify cells. QuPath is an open-source software utilized for digital pathology and image analysis. QuPath has high accuracy with results highly correlated to manual cell counting. This software also processes large datasets quickly and provides an accurate and time-efficient alternative to manual cell counting. High-resolution images of anterior and posterior cortical regions, including the epicenter, hippocampus, thalamus, and ventral midbrain, were acquired at ×20 magnification. Microglial (Iba1+) and astrocytic (GFAP+) cells were identified and quantified using machine learning-based classifiers within QuPath. Automatic cell counting was performed across multiple fields of view per region, and cell density was normalized to the area (cells per mm²). At least three sections per mouse were analyzed, with data compared between anterior and posterior regions post-mTBI.

High-resolution images were analyzed using QuPath software, where machine learning-based classifiers were calibrated by manually annotating a subset of images. This calibration ensured that the automated cell count closely matched manual counting in terms of both accuracy and reproducibility.

### 7.5 Bootstrap Analysis of Group Differences

Given the limited sample size per region (anterior and posterior), conventional parametric statistical tests were not well suited to assess differences between groups. To address this limitation and robustly estimate the uncertainty of our differences, a percentile-based bootstrap resampling analysis for each of the immunohistochemical markers (DAPI, GFAP, Iba1, and NeuN) was conducted. The percentile-based analysis allows for a more comprehensive understanding of the projected outcomes by focusing on the confidence intervals. Bootstrap analysis has been utilized for smaller sample sizes with high accuracy, especially with a conservative approach^20^. Limitations will be further addressed in the conclusion section.

Bootstrap analysis is utilized in preclinical TBI studies, including region-specific responses when the number of available replicates is low. A study analyzing statistical tests in neuroscience with limited sample sizes found that bootstrap tests compared to traditional tests (ANOVA. T-tests, Regression Analysis, etc.), had a lower false positive rate for smaller samples. The bootstrap methods stayed along the threshold along with the SEM. Furthermore, the bootstrap method has a conservative bias for independent datasets^21^. This means that the bootstrap analysis is highly sensitive and ensures robustness. Thus, bootstrap analysis is highly suitable when working with limited samples of raw data.

For each marker, the raw count data is first separated by region.

For each marker, 1,000 bootstrap replicates were generated as follows. The 1,000 benchmark typically provides a stable, reliable estimate of the statistical distribution^18^.

1. **Resampling:** In each replicate, two values were drawn with replacement from the anterior group, and separately from the posterior group to randomly select a value from the anterior dataset, and similarly for the posterior group.
2. **Calculating Replicate Statistics:** For each replicate, the mean of the two resampled anterior values and the mean of the two resampled posterior values were computed. The replicate “difference in means” was then calculated as (Posterior Mean − Anterior Mean) (Table 3 and 4).
3. **Constructing the Bootstrap Distribution:** This procedure was repeated 1,000 times, yielding a distribution of 1,000 differences in means for each stain. This collection provides insights into the quantitative differences that would be expected if additional data were collected under the same conditions.
4. **Determining the 95% Confidence Interval:** The lower and upper bounds of the 95% CI for the difference were obtained by taking, respectively, the 2.5th and 97.5th percentiles of the bootstrap distribution using the percentile function.

**Table 3.** **Percentile-Based Bootstrap Analysis** supporting the qualitative assessment of fluorescence by modeling 1000 replicates per marker and region. The table below summarizes means and 95% confidence intervals.

| Marker | Region | Bootstrap Mean | 95% CI Lower | 95% CI Upper |
| --- | --- | --- | --- | --- |
| DAPI | Posterior | 971 | 848 | 1098 |
|  | Anterior | 1187 | 1035 | 1353 |
| GFAP | Posterior | 115.5 | 113 | 118 |
|  | Anterior | 614 | 485 | 746 |
| Iba1 | Posterior | 65 | 62 | 68 |
|  | Anterior | 198 | 135 | 264 |
| NeuN | Posterior | 189 | 178 | 200 |
|  | Anterior | 163 | 124 | 204 |

*Notes: The “Original Diff.” represents the difference in means calculated directly from the raw data. The table provides statistical validation of IHC analysis with consistent trends of higher inflammation in the anterior region.*

The results of the bootstrap analysis are summarized in Tables 3 and 4 in the results. It should be noted that the bootstrap results are only a projection of a trend or results that are likely to be observed with a larger dataset.

**Table 4.** **Bootstrap Analysis Mean Differences** is a numerical representation and summary of Table 3 to validate the hypothesis of coordinate-specific variation. There were consistently higher levels of activation across stains in the anterior region except for NeuN.

| Marker | Anterior Mean | Posterior Mean | Mean Difference |  |
| --- | --- | --- | --- | --- |
| DAPI | 1187 | 971 | 216 | Higher in Anterior |
| GFAP | 614 | 115.5 | 498.5 | Significantly Higher in Anterior |
| Iba1 | 198 | 65 | 133 | Higher in Anterior |
| NeuN | 163 | 189 | -26 | Slightly Higher in Posterior |

This study used both quantitative and qualitative assessments. The quantitative analysis was conducted through cell quantification and statistical validation as described above. This procedure reiterated results determined from qualitative analysis of fluorescent imagery through cell quantification. The qualitative analysis allowed for the spatial or pattern analysis of microglia and mitochondrial activity. In relation, the quantitative approach determined the coordinate specific and injury-site results.

## 8. Results

### 8.1 Neuroinflammation Varies Across Coordinates (Microglial Analysis)

The inflammatory response to traumatic brain injury (TBI) was evident through differences in the brain morphology following moderate TBI (mTBI) when compared to the sham (control) group. Injured brains displayed visible cavitation at the site of impact, underscoring the structural damage caused by TBI, which was not present in the sham group, ensuring consistency in the procedure. These gross changes highlight the localized nature of the injury and the subsequent inflammatory cascade, as indicated by significant swelling of the brain (Figure 2).

**Figure 2:**
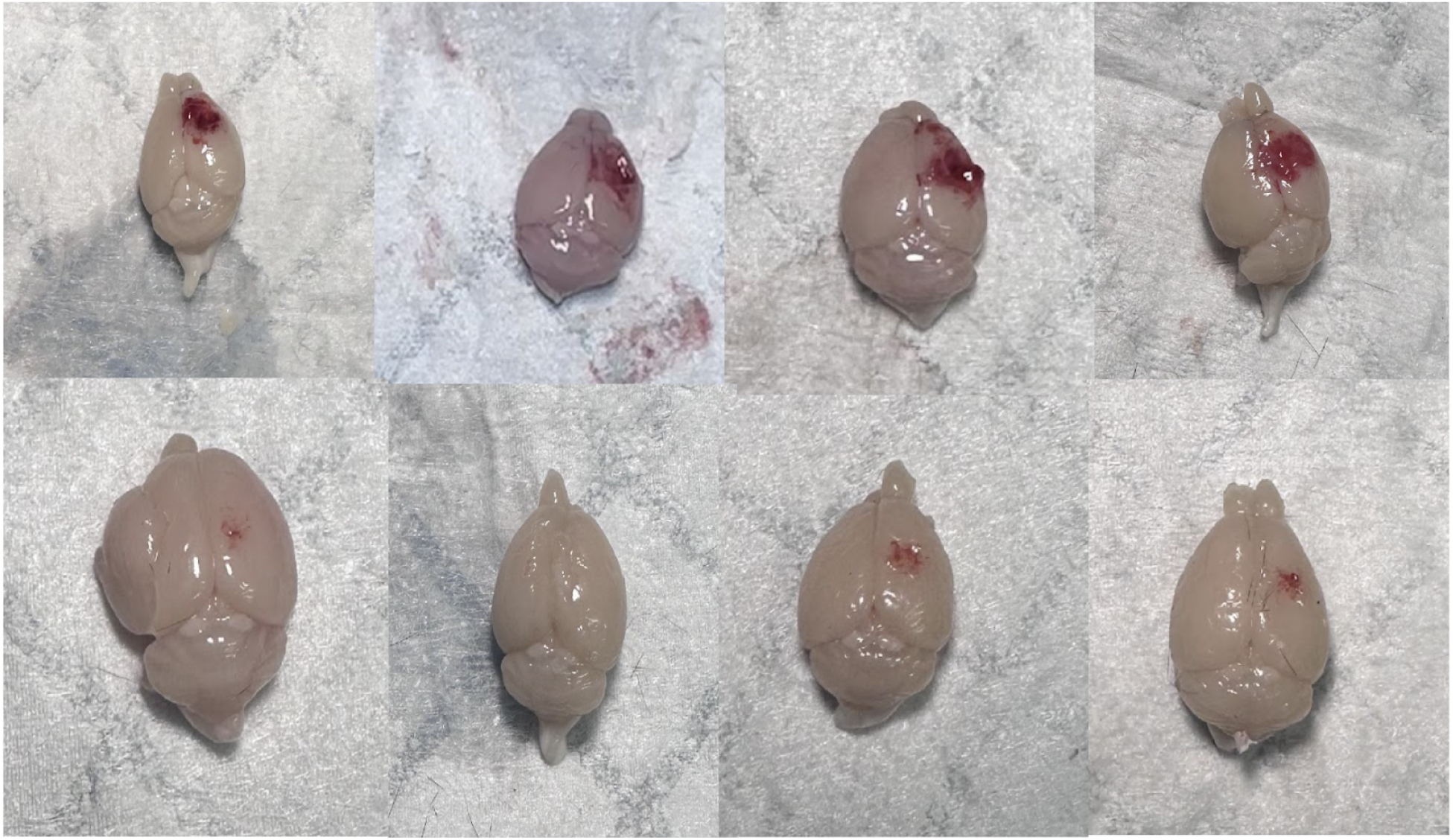
**Gross Anatomy of TBI Samples** are representative of swelling post-injury, confirming the injury and successive inflammation.

Immunohistochemistry (IHC) was conducted to assess the location and extent of the inflammatory response following moderate TBI at both anterior and posterior cortical coordinates. Anterior regions displayed elevated Iba1+ and GFAP+ cell densities and marked neuronal loss (NeuN+ staining), particularly in the hippocampal DG and CA1 regions, compared to the posterior region at 7 DPI (Tables 3 and 4, Figure 4).

NeuN+ staining, which indicates neuronal integrity, showed a disrupted pattern in the anterior injury sites at 7-DPI, consistent with neuronal loss and damage. In contrast, posterior TBI-injured mice exhibited organized NeuN+ staining in these regions. The decrease in NeuN immunoreactivity was most pronounced in the DG and CA1 areas of the hippocampus, with a lesser degree of neuronal loss observed in the CA2 and CA3 areas (Figure 3). immunohistochemical analysis revealed significant differences between anterior and posterior regions following TBI.

**Figure 3:**
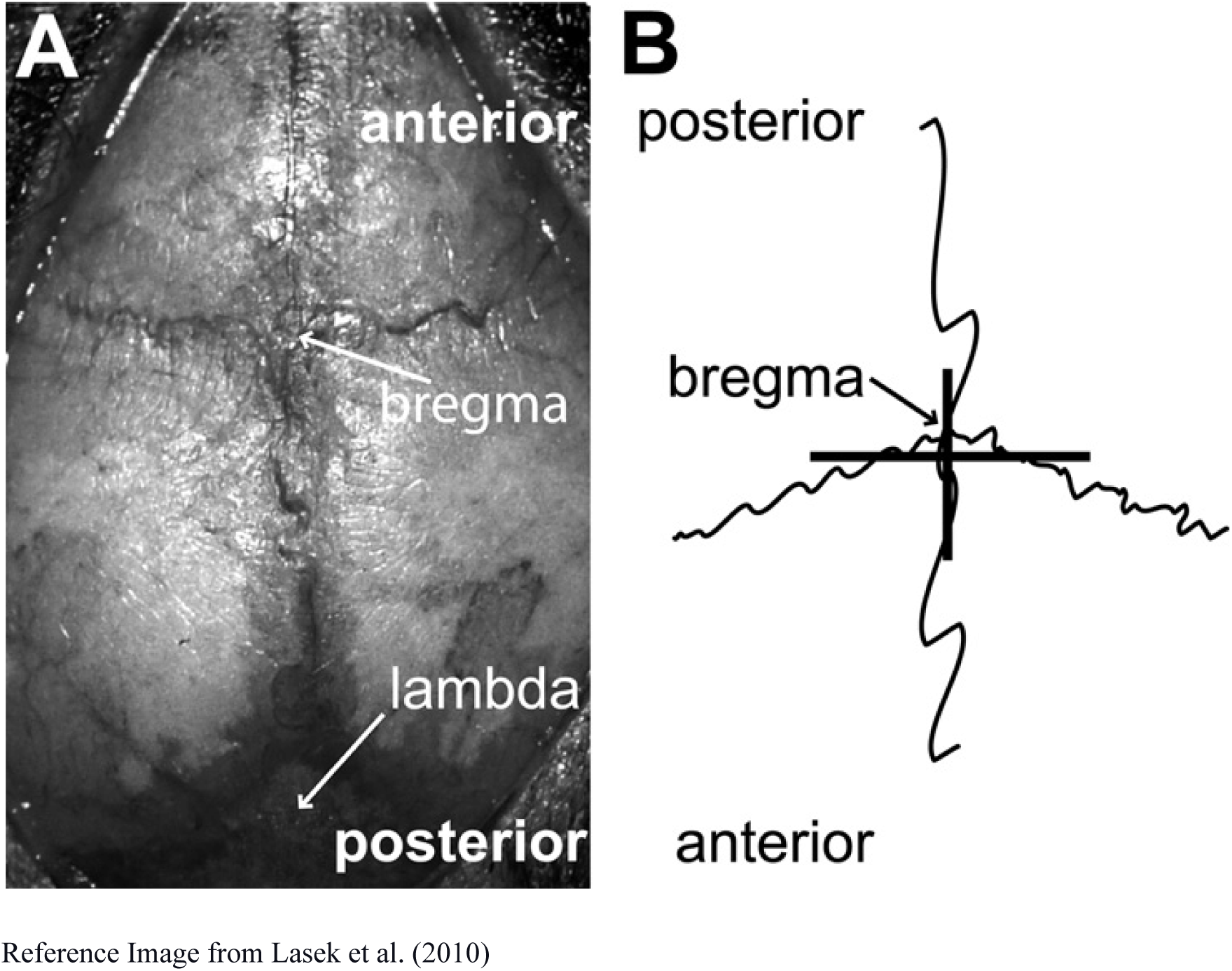
Anterior-Posterior Reference. (A) represents a reference model for the reader’s visualization of the approximated anterior vs posterior coordinates. The points are lined up against the bregma. (B) is the microscope reference of the coordinates.

At the injury site, the increased labeling of Iba1+ and GFAP+ cells at 7-DPI further confirmed the presence of activated microglia and astrocytes, accompanied by a marked reduction in NeuN+ staining, indicating neuronal damage. In the posterior cortical regions, a similar inflammatory pattern was anticipated but was likely more localized to the impact site, with less pronounced effects in remote areas such as the thalamus, ventral midbrain, and corpus callosum.

In Figure 4, the QuPath results from Table 5 are validated by showing clear differences in the posterior and anterior coordinates. These findings are further supported by the results of the bootstrap analysis presented in Tables 3 and 4, as well as in Figure 4. The correlation of qualitative and quantitative results in this study also implicates the validity of utilizing bootstrap tests in TBI studies, considering the limited literature. The confidence intervals for GFAP, Iba1, and NeuN indicate significant variability in astrocytic and microglial activation, underscoring the heightened inflammatory response in anterior regions.

**Figure 4.**
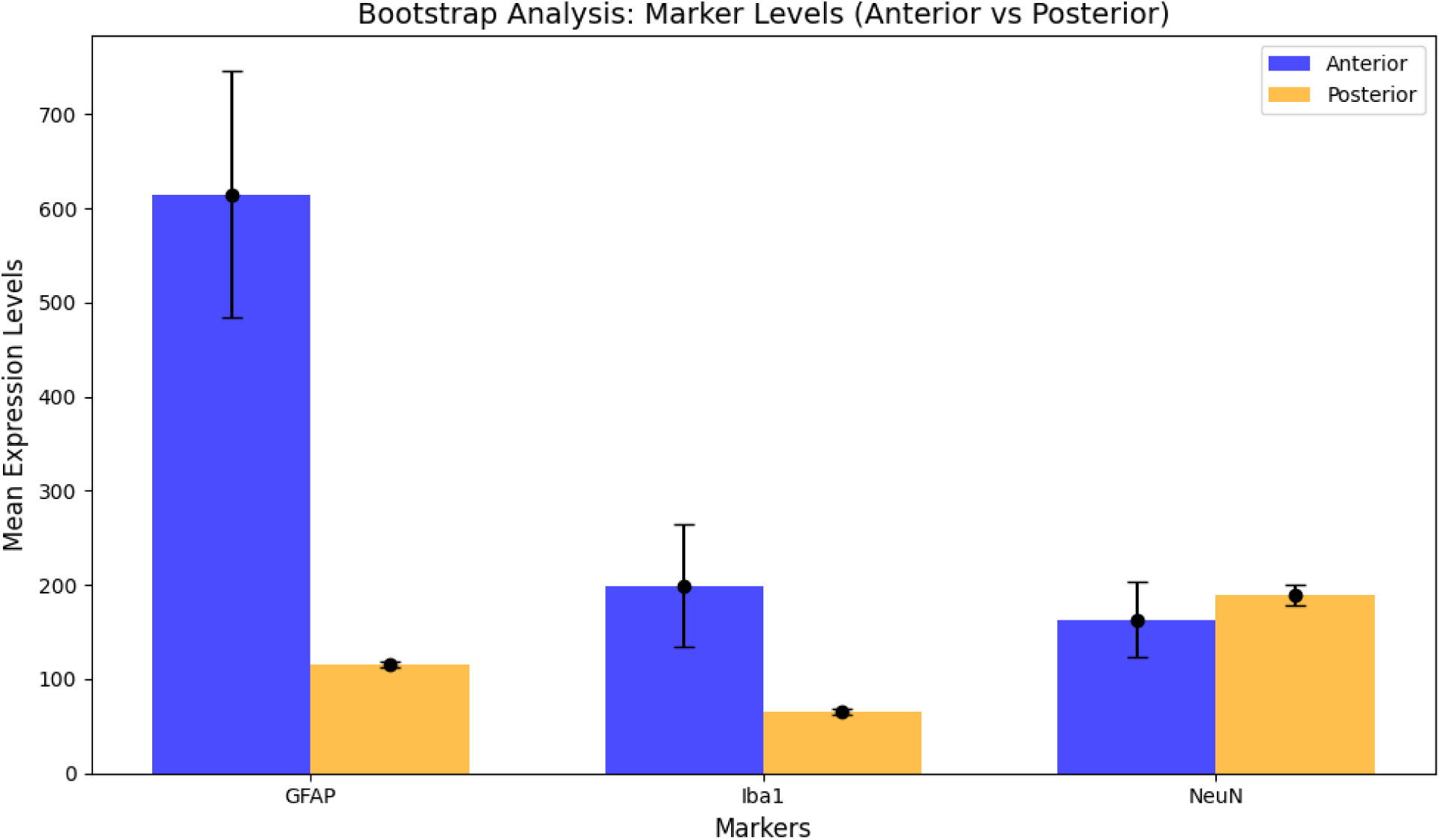
**Bootstrap Analysis** with statistically significant results observed across all stains with the anterior coordinate having higher levels of GFAP, Iba1, and lower levels of NeuN. These results validate IHC results. The figure illustrates anterior versus posterior results with 95% CI per stain.

**Figure 4.**
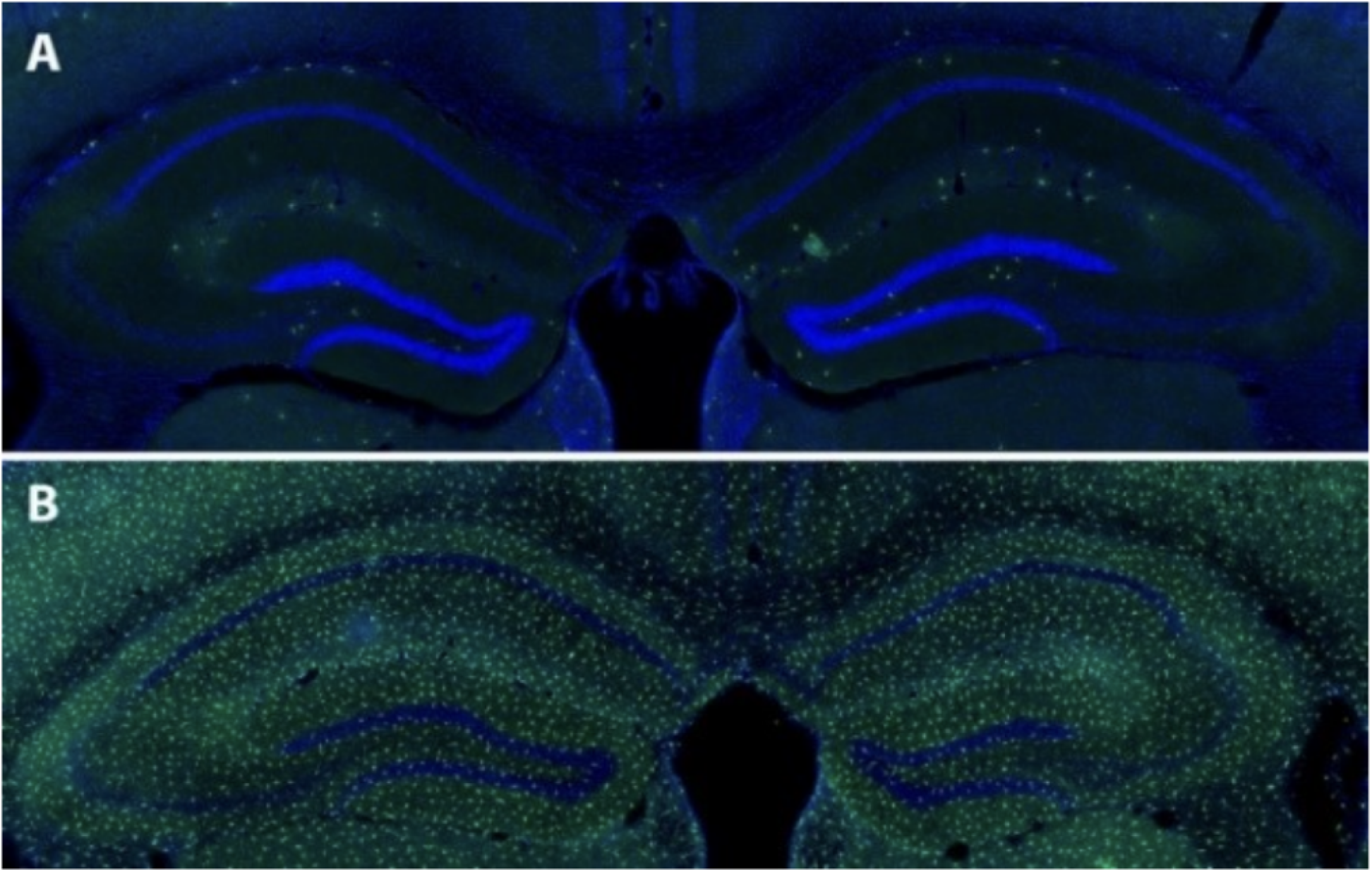
**Immunohistochemistry:** A) Dentate gyrus on sham animal B Dentate gyrus Anterior TBI group. Microglia (green), DAPI (blue). Higher levels of microglial activation and more neuronal death in injury samples as expected due to subsequent inflammation. Figure 4 highlights pronounced microglial activation and neuronal loss in anterior regions, validating the study’s hypothesis.

Starting with the analysis of DAPI, which measures nuclei to reflect cell density, the mean difference of 216 indicates higher levels in the anterior coordinate. The confidence intervals do not overlap, suggesting significant differences in nuclear density post-injury. GFAP levels are markedly elevated in the anterior coordinate, with a mean difference of 498.5; this region also exhibits a wider confidence interval, indicating greater variability in astrocytic activity and supporting the idea of increased activation in the anterior region.

Iba1 levels are significantly higher in the anterior region, with a mean difference of 133. The non-overlapping confidence intervals reinforce this statistical significance and point to localized inflammation. The posterior regions displayed greater neuronal integrity, as indicated by higher NeuN staining values in Table 4. This indicates a moderate decrease in neuron numbers, with marginally better neuronal preservation compared to the anterior coordinate. Statistical comparisons of microglia reveal more dramatic activation in the anterior region compared to the posterior, aligning with qualitative analyses from immunofluorescence studies.

Although both the anterior and posterior coordinates of mTBIs were conducted under the same conditions—using an impactor probe velocity of 3 m/s, with the same dwell time and impactor length—the anterior coordinate showed a significantly greater increase in inflammation, neuronal damage, and activated microglia compared to the posterior region. This suggests that the posterior region of the brain may be deeper or require a greater impact to elicit similar effects. Additionally, there may be variations in microglial density or differences in the prominence of pro-inflammatory cytokines.

Overall, these results highlight inherent differences in brain pathophysiology that must be considered when developing therapeutics. They also suggest that the characterization of moderate TBI in animal models should vary based on the location of the injury, as the indicators of moderate TBI in the anterior region do not directly correlate with the cellular changes observed in the posterior region.

### 8.2 Mitochondrial Dysfunction and Correlating TBI Conditions

The following mitochondrial analysis is based on a qualitative assessment of immunofluorescence. Existing literature has explored the region-specific events associated with mitochondrial dysfunction. An analysis of therapeutic strategies following traumatic brain injury (TBI) found that oxidative damage was more pronounced in the CA3 region compared to the CA1 region of the hippocampus^22^. This finding correlates with a comparative assessment of inflammation conducted across anterior versus posterior approaches. However, there remains a gap in understanding mitochondrial behavior using function-specific stains, particularly in the context of a deeper angle impact, as illustrated by this study’s reproducible method of controlled cortical impact (CCI)^17^. Therefore, we evaluated the effectiveness of three stains—MitoGreen, TOM20, and MFN1—to gain insights into mitochondrial dysfunction.

MitoGreen is a commonly used fluorescent stain for visualizing mitochondria. However, it primarily provides general mitochondrial visualization based on mass or abundance rather than reflecting their functional state^8^. Our analysis using MitoGreen yielded inconclusive results. We then turned our attention to TOM20, which is part of the outer mitochondrial membrane complex and plays a role in importing proteins into mitochondria. While TOM20 is useful for visualizing mitochondrial abundance and distribution, it does not offer insights into mitochondrial dysfunction^5^. In contrast, MFN1, which is directly involved in the fusion process of mitochondria, can provide valuable information regarding mitochondrial function and aligns with the fifth aim of this study. Unlike MitoGreen and TOM20, MFN1 offers a unique advantage in assessing mitochondrial dysfunction by highlighting fusion processes critical to energy metabolism and repair mechanisms post-injury^29^. There is also a need to address gaps in using MFN1 as a mitochondrial stain to uncover its therapeutic potential for enhancing mitochondrial fusion and enable more accurate identification of mitochondrial abnormalities^29^.

The MFN1 stains revealed mitochondrial activity through fluorescence, as shown in Image A (Figure 5). This activation corresponds with the increased activity of astrocytes (Figure 5.C) and the lack of neuronal activity (Figure 5.B) at the injury site. The persistent mitochondrial activity observed, despite observing the stain of 7 DPI, indicates ongoing repair mechanisms under inflammatory conditions. Additionally, the lower intensity of mitochondrial fluorescence compared to astrocyte and neuronal activity suggests some degree of mitochondrial dysfunction. This weaker fluorescence may be due to diminished mitochondrial activity in neuronal cells, as indicated by the absence of NeuN fluorescence (Figure 5.B). The existing fluorescent signal could indicate astrocytic mitochondria working to counter neuroinflammation following the injury.

**Figure 5.**
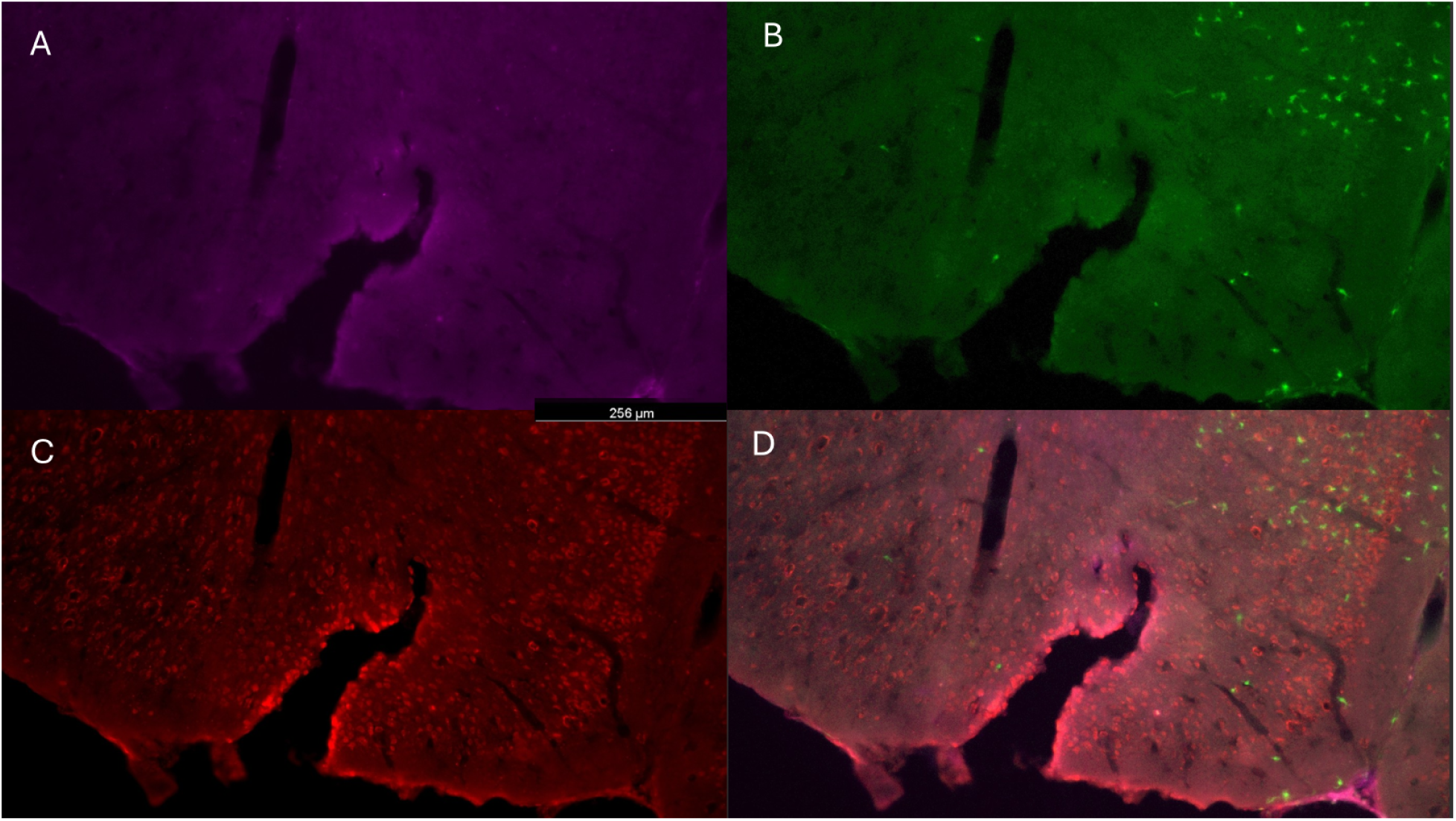
**Mitochondrial IHC** reveals A (MFN1), B (NeuN), C (GFAP), D (overlay). Mitochondrial stains are posterior-TBI. MFN1 staining reveals mitochondrial activity at injury sites. Panels highlight increased astrocyte activity (GFAP) and diminished neuronal fluorescence (NeuN), illustrating persistent inflammation and mitochondrial dysfunction.

The observed variability in inflammatory and neuronal responses across brain regions highlights critical insights into TBI pathophysiology, which will be explored in the discussion section.

## 9. Conclusion

This study investigated the spatial distribution of the inflammatory response and neuronal damage following moderate traumatic brain injury (TBI) at both anterior and posterior cortical coordinates. The findings reveal significant differences in the inflammatory and neuronal responses between these two brain regions, particularly regarding microglial activation and astrocyte response.

The anterior brain regions demonstrated significant disruption in the organization of NeuN+ cells, reflecting neuronal loss and potential synaptic dysfunction. These findings support previous studies suggesting that areas closer to the impact site experience more severe neuroinflammation and neuronal loss following TBI^27^. The sustained activation of microglia and astrocytes at the injury site indicates that neuroinflammatory processes continue well beyond the initial injury, potentially contributing to long-term neurodegenerative outcomes, including cognitive impairments. The anterior region’s heightened neuronal damage may be attributed to differences in neuronal subtypes that are more prone to excitotoxicity and neuroinflammatory cascades post-injury.

In summary, this study provides compelling evidence of regional differences in inflammatory and neuronal responses to moderate TBI. The anterior cortical regions show significantly greater microglial activation, astrocyte proliferation, and neuronal damage compared to the posterior regions. These findings underscore the importance of considering region-specific effects when studying TBI pathophysiology and developing targeted interventions^14^. Region-specific treatments, such as anti-inflammatory drugs or neuroprotective agents, could be tailored to the anterior regions to mitigate damage following mTBI. Anterior-focused therapies such as imaging techniques might incorporate cognitive exercises to rebuild neuronal connections in areas with high inflammation and synaptic dysfunction.

This study also highlights the significance of qualitative immunofluorescence analysis in understanding mitochondrial dynamics post-TBI. Consistent with previous findings, oxidative damage was more pronounced in the CA3 region compared to the CA1 of the hippocampus, emphasizing region-specific vulnerability^22^. Furthermore, this analysis connects existing knowledge on anterior versus posterior inflammation with functional studies, revealing a critical gap in assessing mitochondrial behavior using function-specific stains in deeper impact models.

Our evaluation of mitochondrial stains—MitoGreen, TOM20, and MFN1—demonstrated that MFN1 provides more insightful data on mitochondrial dysfunction. MFN1 stains correlated mitochondrial activity with astrocyte-driven repair mechanisms under inflammatory conditions. This suggests the role of astrocytic mitochondria in mitigating neuroinflammation following TBI.

The weaker fluorescence observed in neurons, attributed to the absence of NeuN activity, underscores persistent mitochondrial dysfunction. These findings reinforce the need for targeted research on MFN1 as a mitochondrial stain, not only to elucidate its role in post-TBI repair but also to explore its therapeutic potential in enhancing mitochondrial fusion and recovery mechanisms.

Having established coordinate-specific variability in inflammation and neuronal damage, this study explores the broader implications for therapeutic development.

### 9.1 Limitations and Future Directions

This study has several limitations. First, there is no widely accepted scoring system for assessing injury severity in animals, comparable to the Glasgow Coma Scale (GCS) used for human patients with traumatic brain injury (TBI). Moreover, scoring systems can vary across laboratories, which, along with minor differences in craniotomy procedures or cell loss, complicates the comparison of findings from different studies^27^. The discrepancies observed in this study regarding inflammatory responses at two brain coordinates provide insights for developing a severity scale for animal models of TBI. It is assumed that injuries closer to the anterior region, where a more profound injury response was noted, would be indexed as higher severity on this scale. The establishment of a standardized protocol for moderate TBI induction using controlled cortical impact (CCI) offers a reproducible method for future studies. Bootstrap analysis of additional replicates, as validated through this study, could aid in creating this severity index. Researchers can apply this protocol to explore deeper impacts and region-specific responses, fostering more translational animal models.

Additionally, no ANOVA or t-tests were performed for further statistical analysis due to the limited sample size. Instead, this study employed bootstrap analysis to determine numerical significance. As described in the results section, the QuPath cell quantification and the subsequent transfer of this data to a graph (Figure 5) support the findings by following methods from previous studies with similar sample sizes^23^. However, the bootstrap analysis was also constrained by the small sample size. Future studies with larger sample sizes across various coordinates could draw definite patterns and trends from the raw data presented in Appendix B.

Another area for future research includes analyzing trends in MFN1 within the anterior region to replicate the comparative assessment of microglial activation discussed in this study^13^. Additionally, further studies should explore mitochondrial function separate from neuronal death and astrocyte activity to achieve more conclusive insights into abnormal energy metabolism. These additional findings can build on the reproducible method of controlled cortical impact (CCI) for moderate TBI presented in this research. Enhancing mitochondrial fusion through MFN1-targeted treatments could mitigate energy deficits and promote neuronal survival, offering new avenues for post-TBI recovery.

Future research should also consider regional variations in analyzing how mitochondrial dysfunction and pro-inflammatory microglial activation are affected following microglial depletion. To date, only holistic studies have examined microglial depletion in chronic TBI to mitigate neurodegeneration and alleviate neurological deficits^1^. The focus on short-term depletion is particularly relevant, given the critical roles microglia play in restoring tissue homeostasis after a pathological insult. For instance, treatment with a CSF1R inhibitor demonstrated that even when administered four months post-injury, it significantly reduced neuronal damage and dysfunction ^9^. A potential avenue of study would involve applying the coordinate variation patterns illustrated in this research while implementing microglial depletion. The marked increase in microglial activation in the anterior compared to the posterior coordinates suggests the need to adjust the intensity of microglial depletion for region-specificity.

Further examination of microglia is necessary due to their therapeutic potential having shown mixed results in various studies. While microglia depletion has been associated with reduced neuronal damage^9^, another study highlighted the neuroprotective role of microglia in stroke^24^. Microglia have also been implicated in angiogenesis, the process through which new blood vessels develop from existing ones. However, eliminating microglia has been linked to a decline in the retinal arrangement of blood vessels. Stroke can cause synaptic dysfunction, but both microglia and macrophages play crucial roles in synaptic remodeling. Furthermore, regulating the activation states of microglia and macrophages could alter the inflammatory environment following stroke, encouraging neuroprotection and angiogenesis^24^. Due to the contrasting findings of these studies, there is a pressing need for more research into the role of microglia in inflammatory pathways. Gaining a deeper understanding of these pathways is vital for investigating the interactions between astrocyte activation, microglia, and mitochondrial activity.

## Appendix A: List of Abbreviations

ANOVA: Analysis of Variance
a-syn: Alpha synuclein
ATP: Adenosine Triphosphate
CA: Cornu Ammonis
CCI: Controlled Cortical Impact
CSF1R: Colony-stimulating factor 1 receptor
DG: Dentate Gyrus
DNA: Deoxyribonucleic Acid
DPI: Days Post-injury
GFAP: Glial Fibrillary Acidic Protein
IACUC: Institutional Animal Care and Use Committee
Iba1: Ionized calcium binding adaptor molecule 1
IHC: Immunohistochemistry
LPS: Lipopolysaccharide
M1: Type 1 Microglia
M2: Type 2 Microglia
MFN1: Mitofusin 1
mTBI: Moderate Traumatic Brain Injury
NeuN: Neuronal nuclear protein
OCT: Optimal Cutting Temperature compound
PBS: Phosphate-buffered Saline
PFA: Paraformaldehyde
ROS: Reactive Oxygen Species
TBI: Traumatic Brain Injury
TOM20: Translocase of Outer Mitochondrial Membrane 20

## Appendix B: Quantitative Microglial Density and Activation

Reference 1. Bootstrap Replicate Results

**Table B1.**
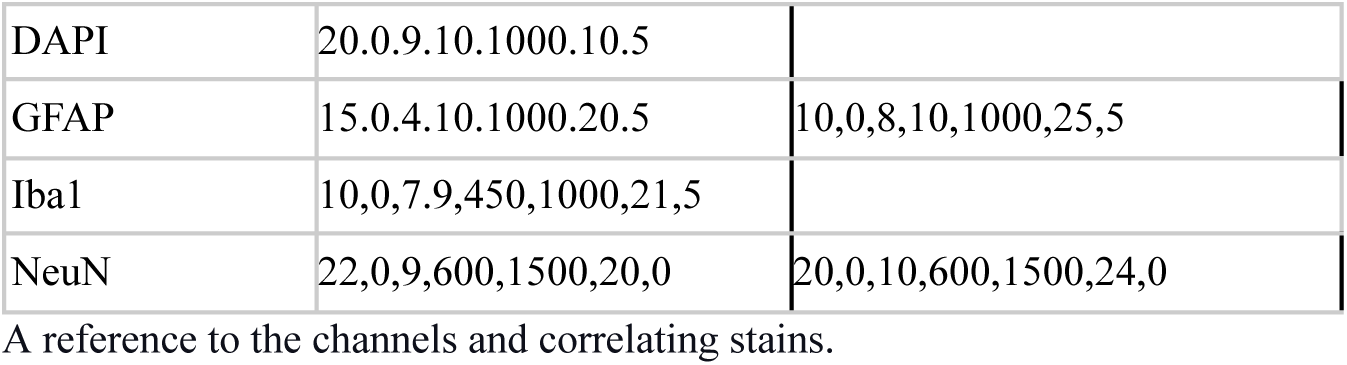
Cell Detection Channels

|  |  |  |
| --- | --- | --- |
| DAPI | 20.0.9.10.1000.10.5 |  |
| GFAP | 15.0.4.10.1000.20.5 | 10,0,8,10,1000,25,5 |
| Iba1 | 10,0,7.9,450,1000,21,5 |  |
| NeuN | 22,0,9,600,1500,20,0 | 20,0,10,600,1500,24,0 |

A reference to the channels and correlating stains.

**Table B2.** Coordinate-specific IHC Quantification

| Region | Sample | DAPI | GFAP | Iba1 | NeuN |
| --- | --- | --- | --- | --- | --- |
| Posterior | MN143_18_s4 | 1126 | 118 | 63 | 182 |
| Posterior | MN2_18_s5 | 816 | 113 | 67 | 196 |
| Anterior | MN3_18_s4 | 899 | 307 | 55 | 241 |
| Anterior | MN4_18_s5 | 1475 | 921 | 341 | 85 |

IHC cell quantification results in correlation to coordinate-specific samples.

**Table B3.** Standard Bootstrap Analysis

| Marker | Region | Mean | Median | StdDev | Range | Bootstrap Difference Mean | 95% CI (Lower) | 95% CI (Upper) |
| --- | --- | --- | --- | --- | --- | --- | --- | --- |
| DAPI | Anterior | 1187 | 1187 | 288 | 576 | -214.08 | -659 | 227 |
|  | Posterior | 971 | 971 | 155 | 310 |  |  |  |
| GFAP | Anterior | 614 | 614 | 307 | 614 | -497.93 | -808 | -189 |
|  | Posterior | 115.5 | 115.5 | 2.5 | 5 |  |  |  |
| Iba1 | Anterior | 198 | 198 | 143 | 286 | -131.25 | -278 | 12 |
|  | Posterior | 65 | 65 | 2 | 4 |  |  |  |
| NeuN | Anterior | 163 | 163 | 78 | 156 | 26.41 | -59 | 111 |
|  | Posterior | 189 | 189 | 7 | 14 |  |  |  |

**Table B4.** Percentile-based Bootstrap Analysis

| Marker | Region | Bootstrap Mean | 95% CI Lower | 95% CI Upper |
| --- | --- | --- | --- | --- |
| DAPI | Posterior | 971 | 848 | 1098 |
|  | Anterior | 1187 | 1035 | 1353 |

\*\*Abbreviations in Appendix A

|  |  |  |  |  |
| --- | --- | --- | --- | --- |
| GFAP | Posterior | 115.5 | 113 | 118 |
|  | Anterior | 614 | 485 | 746 |
| Iba1 | Posterior | 65 | 62 | 68 |
|  | Anterior | 198 | 135 | 264 |
| NeuN | Posterior | 189 | 178 | 200 |
|  | Anterior | 163 | 124 | 204 |

## Notes

### Competing Interest Statement

The authors have declared no competing interest.

